# White matter hyperintensities are common in midlife and already associated with cognitive decline

**DOI:** 10.1101/687111

**Authors:** Tracy d’Arbeloff, Maxwell L. Elliott, Annchen R. Knodt, Tracy R. Melzer, Ross Keenan, David Ireland, Sandhya Ramrakha, Richie Poulton, Tim Anderson, Avshalom Caspi, Terrie E. Moffitt, Ahmad R. Hariri

**Author notes:** Correspondence: Ahmad R. Hariri, Ph.D., Professor of Psychology and Neuroscience Director, Laboratory of NeuroGenetics Head, Cognition and Cognitive Neuroscience Training Program, Duke University Durham, NC 27708, USA. These authors contributed equally to this work (Tracy d’Arbeloff, Maxwell L. Elliott). Abbreviations: ADRD - Alzheimer’s disease and related dementias, BMI – body mass index, WMH – white matter hyperintensity, FLAIR – fluid attenuated inversion recovery.

## Abstract

White matter hyperintensities (WMHs) proliferate as the brain ages and are associated with increased risk for cognitive decline as well as Alzheimer’s disease and related dementias. As such, WMHs have been targeted as a surrogate biomarker in intervention trials with older adults. However, it is unclear at what stage of aging WMHs begin to relate to cognition and if they may be a viable target for early prevention. In a population-representative birth cohort of 843 45-year-olds we measured WMHs using T2-weighted MRI, and we assessed cognitive decline from childhood to midlife. We found that WMHs were common at age 45 and that WMH volume was modestly associated with both lower childhood (ß=-0.08, *p*=0.013) and adult IQ (ß=-0.15, *p*<0.001). Moreover, WMH volume was associated with greater cognitive decline from childhood to midlife (ß=-0.09, *p*<0.001). Our results demonstrate that a link between WMHs and early signs of cognitive decline is detectable decades before clinical symptoms of dementia emerge. Thus, WMHs may be a useful surrogate biomarker for identifying individuals in midlife at risk for future accelerated cognitive decline and selecting participants for dementia prevention trials.

## Introduction

To address rising economic and health burdens due to Alzheimer’s disease and related dementias (ADRD), government funding for aging research has more than quadrupled in recent years (Kaiser, 2018). However, the success of this investment hinges on developing surrogate biomarkers—biological measures that are part of the putative disease pathway and are measurable before the onset of clinical symptoms—so that prevention can target at-risk individuals before cerebral decline has taken hold. Successful surrogate biomarkers would allow clinicians to assess risk, monitor sub-clinical disease progression, and intervene before clinically significant dementia symptoms manifest.

Research shows white matter hyperintensities (WMHs) are one such surrogate biomarker of cognitive decline and ADRD that can be measured in the brains of older adults (Cees De Groot *et al.*, 2000; Lee *et al.*, 2016). As the brain ages, it begins to accrue small microbleeds and lesions in white matter that are detectable as WMHs using fluid-attenuated inversion recovery (FLAIR) magnetic resonance imaging (MRI) (Iadecola, 2013). While WMHs are uncommon in adults before age 30 (Habes *et al.*, 2016), they are detectable in over 90% of individuals by age 65 (Longstreth *et al.*, 1996). In older adults, WMHs are associated with multiple dementia risk factors, including increasing age, hypertension, stroke, brain atrophy, and cognitive ability (Prins and Scheltens, 2015). Longitudinal studies in older adults have reported that the spread of WMHs contributes to elevated risk for ADRD and coincides with age-related cognitive decline (Debette and Markus, 2010). Furthermore, baseline WMH load at mean age 62 can predict the onset of dementia up to 20 years later (Coker *et al.*, 2019).

Consequently, WMHs have been targeted as a surrogate biomarker for dementia prevention trials (Debette and Markus, 2010). However, these trials have produced mixed results (Prins and Scheltens, 2015). A limitation of existing trials is that they have targeted older adults in their 60s, 70s and 80s. Older brains are characterized by age-related deterioration and may be less responsive to intervention (Sperling *et al.*, 2014; Moffitt *et al.*, 2017). One solution is to assess WMHs in midlife, a time when the brain may be more responsive to interventions and has yet to be affected by decades of age-related organ decline. It is known that WMHs predict cognitive decline and risk for ADRD in older adults (Valdés Hernández *et al.*, 2013), but it is not known when WMHs accumulate sufficiently to be associated with early cognitive decline.

Here, we tested the hypotheses that WMHs are detectable in midlife and already associated with cognitive decline from childhood in a population-representative birth cohort aged 45 years. Support for these hypotheses would provide novel evidence that WMHs could be a surrogate biomarker of risk in the general population as early as midlife, allowing for earlier—and potentially more effective—interventions for cognitive decline and ADRD.

## Materials and methods

### Study design and population

Participants were members of the Dunedin Multidisciplinary Health and Development Study, a longitudinal investigation of health and behavior in a population representative birth cohort. The full cohort (N=1,037; 91% of eligible births; 52% male) comprises all individuals born between April 1972 and March 1973 in Dunedin, New Zealand, who were eligible based on residence in the province and who participated in the first assessment at age 3 years. The cohort represents the full range of socioeconomic status in the general population of New Zealand’s South Island (Poulton *et al.*, 2015). The cohort matches the New Zealand National Health and Nutrition Survey on adult health indicators (e.g., BMI, smoking, primary-care visits) and the NZ Census on educational attainment. The cohort is primarily white (93%), which matches the demographics of the South Island (Poulton *et al.*, 2015). Assessments were carried out at birth and ages 3, 5, 7, 9, 11, 13, 15, 18, 21, 26, 32, 38, and most recently (completed April 2019) 45 years, when 94.1% (N=938) of the 997 participants still alive took part. 875 (93% of age-45 participants) also completed MRI scanning. Scanned participants did not differ from other living participants on childhood SES or childhood IQ (see attrition analysis in the **Supplementary Material**). The relevant ethics committees approved each phase of the Study and informed consent was obtained from all participants.

### Measurement of cognitive ability

Cognitive ability in adulthood was assessed using the Wechsler Adult Intelligence Scale–IV (WAIS-IV; (IQ score range, 40 - 160) at age 45 years (Weschler, 2008). Cognitive ability in childhood was assessed using the Wechsler Intelligence Scale for Children Revised (WISC-R; score range, 40 - 160) at ages 7, 9, and 11 years with the mean for these three assessments used in analyses (Wechsler, 1974; Moffitt *et al.*, 1993).

### Imaging parameters

Each participant was scanned using a Siemens Skyra 3T scanner equipped with a 64-channel head/neck coil at the Pacific Radiology imaging center in Dunedin, New Zealand. High resolution structural images were obtained using a T1-weighted MP-RAGE sequence with the following parameters: TR = 2400 ms; TE = 1.98 ms; 208 sagittal slices; flip angle, 9°; FOV, 224 mm; matrix =256×256; slice thickness = 0.9 mm with no gap (voxel size 0.9×0.875×0.875 mm); and total scan time = 6 min and 52 s. 3D fluid-attenuated inversion recovery (FLAIR) images were obtained with the following parameters: TR = 8000 ms; TE = 399 ms; 160 sagittal slices; FOV = 240 mm; matrix = 232×256; slice thickness = 1.2 mm (voxel size 0.9×0.9×1.2 mm); and total scan time = 5 min and 38 s.

### Quantification of white matter hyperintensities

To identify and extract WMH volume, T1-weighted and FLAIR images for each participant were run through UBO Detector (Jiang *et al.*, 2018), a cluster-based, fully-automated, pipeline that uses FMRIB’s Automated Segmentation Tool (Zhang *et al.*, 2001) to identify candidate clusters. Using K-nearest neighbors (k-NN) algorithms, clusters in the MRI images are classified as WMHs or non-WMHs (i.e., grey matter or cerebral spinal fluid) based on anatomical location, intensity, and cluster size features. A Diffeomorphic Anatomical Registration through Exponentiated Lie (DARTEL) template of 55 years or younger was used to best approximate the age of our cohort (Ashburner, 2007), and a grey matter mask was applied to decrease the chance of false positives. The resulting WMH probability maps were thresholded at 0.7, which is the suggested standard (Jiang *et al.*, 2018).

We chose the UBO pipeline because of its high reliability in our data (test-retest ICC = 0.87) and its out-of-sample performance (Jiang *et al.*, 2018). However, for additional quality assurance, every participant’s UBO-generated WMH map was visually inspected to check for false positives (e.g., areas such as the septum that appear similar to WMHs on FLAIR images). Of the 875 scanned participants who had at least 1 MRI scan, 867 had both a T1 and a FLAIR image that are required to extract WMHs with UBO. 843 participants were included in the final analysis after 8 participants were removed for excessive UBO false positives and 4 were excluded because they had incidental findings that interfered with the UBO algorithm, 3 were removed for having multiple sclerosis and 9 participants were excluded for missing IQ data in childhood or adulthood.

### Statistical analyses

All statistical analyses were done using R (v.3.4.5). First, descriptive statistics were generated for the sample as a whole (Table 1). Second, WMH volume was log transformed for normality. Third, the associations between WMH volume (measured in mm^3^) and adult IQ and between volume and childhood IQ were tested using ordinary least squares multiple regression. Fourth, the association between volume and change in IQ was tested using ordinary least squares multiple regression. To calculate residualized change scores, volume was regressed on adult IQ, adjusting for childhood IQ. Sex and total brain volume were used as covariates in all analyses.

**Table 1.** Demographic characteristics for the 843 participants from the Dunedin Study included in the current analyses.

|  | **Total**<br>**(n= 843)** | **Men**<br>**(n= 428)** | **Women**<br>**(n=415)** |
| --- | --- | --- | --- |
| **Total Brain Volume (cm<sup>3</sup>)** | 1159.00 ± 117.55 | 1229.55 ± 98.09 | 1086.08 ± 87.76 |
| **BMI (kg/m<sup>2</sup>)** | 28.53 ± 5.81 | 28.54 ± 4.73 | 28.51 ± 6.74 |
| **HbA1 IFCC (US style %)** | 5.67 ± .54 | 5.75 ± .58 | 5.62 ± .48 |
| **Cholesterol (mmol/L)** | 5.15 ± .97 | 5.33 ± 1.01 | 4.97 ± .93 |
| **Systolic BP (mmHg)** | 121.32 ± 14.62 | 125.46 ± 13.88 | 117.02 ± 14.14 |
| **Diastolic BP (mmHg)** | 80.29 ± 10.21 | 84.43 ± 9.40 | 76.01 ± 9.21 |
| **On BP medication (n)** | 59 | 31 | 28 |
| **On Cholesterol medication (n)** | 32 | 25 | 7 |
| **On Medication for Heart Problems (n)** | 9 | 6 | 4 |
| **Heart Attack (has/had) (n)** | 6 | 5 | 1 |
| **Afib (has/had) (n)** | 4 | 2 | 2 |
| **Cardiomyopathy (has/had) (n)** | 1 | 1 | 0 |
| **Blocked Arteries (has/had) (n)** | 2 | 2 | 0 |
| **Current Smokers (yes/no) (n)** | 176 | 95 | 81 |
| **Education Attainment (%)** |  |  |  |
| *% No qualification* | 14.6% | 17.8% | 11.3% |

The premise and analysis plan for this project were pre-registered on https://sites.google.com/site/dunedineriskconceptpapers/documents. Analyses reported here were checked for reproducibility by an independent data-analyst, who recreated the code by working from the manuscript and applied it to a fresh dataset.

### Data availability

The dataset reported in the current article is not publicly available due to lack of informed consent and ethical approval but is available from the corresponding author on reasonable request by qualified scientists. Requests require a concept paper describing the purpose of data access, ethical approval at the applicants’ university, and provision for secure data access. Details are available at https://sites.google.com/site/dunedineriskconceptpapers/documents.

## Results

WMHs were common in the cohort, with an average volume of 953.50 mm^3^ (25^th^-75^th^ quartile = 425.25 - 1,142.44 mm^3^, median = 681.75; Fig 1). WMHs were most common around the anterior and posterior horns of the lateral ventricles (e.g., Fig 2).

**Figure 1.**
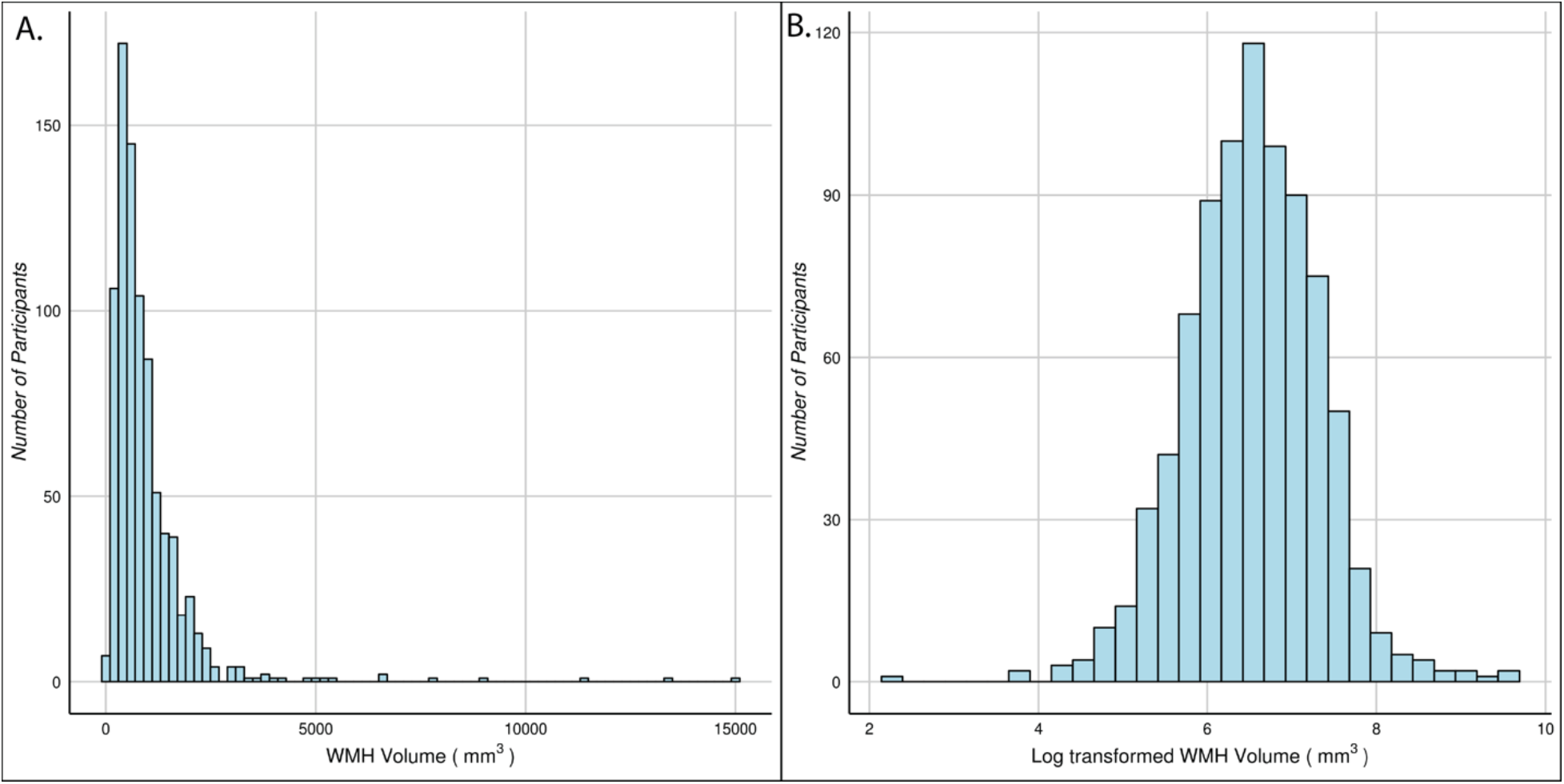
WMHs were common in 45-year-old participants from the Dunedin Study. Panel A shows the distribution of the raw WMH volumes. Panel B shows the log transformation of the volume distribution in panel A. All analyses reported used log-transformed volume.

**Figure 2.**
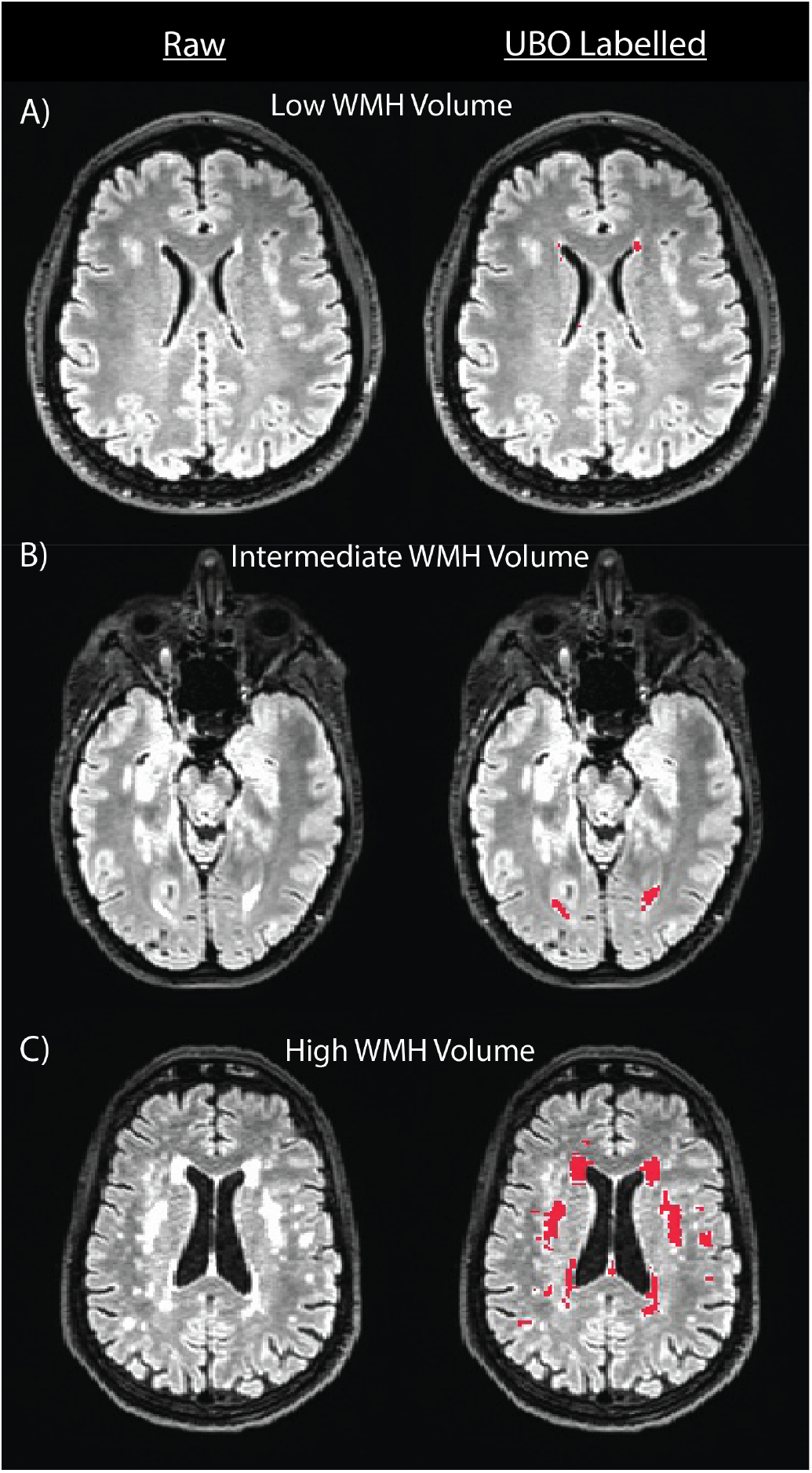
Images depicting relatively low, intermediate, and high WMH-load participants from the Dunedin Study. The left column presents a raw FLAIR image for three representative participants with low, intermediate, and high WMH load. The right column presents UBO labeling (red) of WMHs in the raw images from the left column. As can been seen in these images, WMHs were most common around the anterior and posterior horns of the lateral ventricles as expected. Note that UBO labeling in septal regions was removed from estimation of WMH volume using an exclusion mask.

Larger WMH volume was associated with lower IQ in childhood (ß = −0.08, 95% CI = - 0.15 to −0.02; *p* = 0.013; Fig 3a); individuals with the highest volume (> 1.5 SDs above the mean) had childhood IQs that were 4.80 points lower on average than individuals with the lowest volume (< 1.5 SDs below the mean). This difference was exacerbated in adulthood; larger WMH volume was associated with lower IQ (ß = −0.15, 95% CI = −0.22 to −0.09; *p* < 0.001; Fig 3b) and individuals with the highest volume had adult IQs that were 8.91 points lower than the those with the lowest volume. Lastly, participants with larger WMH volume experienced more cognitive decline by midlife (ß = −0.09, 95% CI = −0.13 to −0.02; *p* < 0.001; Fig 3c).

**Figure 3.**
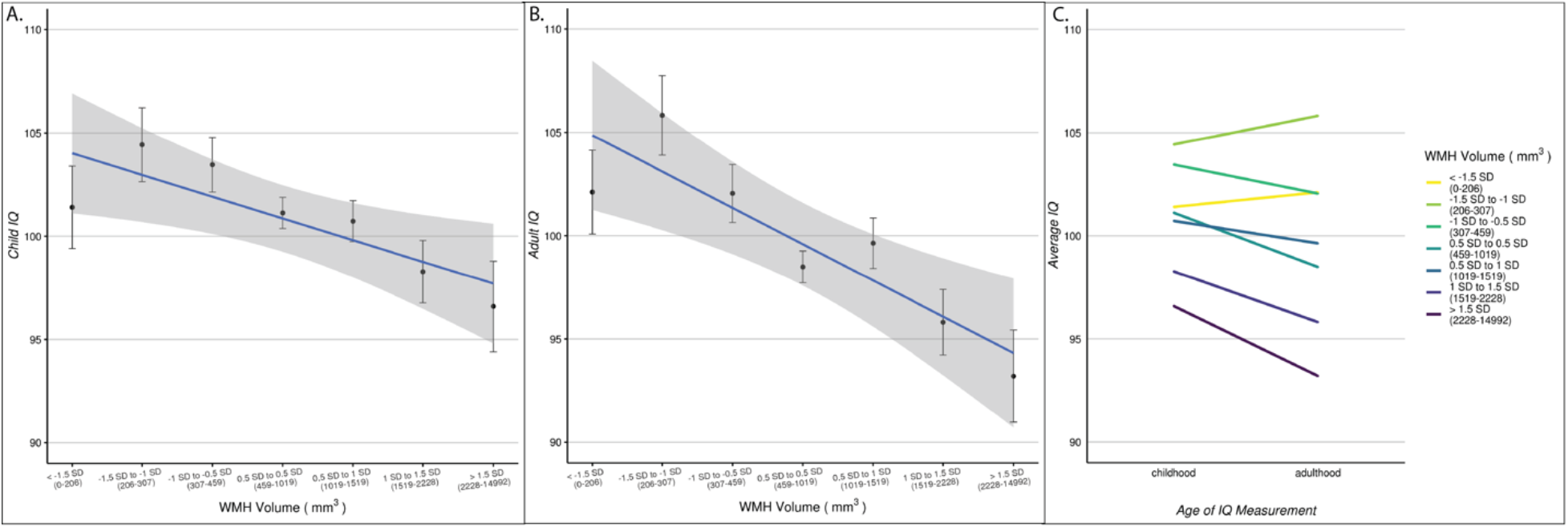
WMHs were associated with cognitive ability and cognitive decline in 843 45-year-old participants from the Dunedin Study. WMHs in all panels are defined by standard deviations (SDs) from the mean volume (mm^3^) ranging from −1.5 to 1.5 SDs in 0.5 SD increments. Sample sizes for each group from the lowest to the highest WMH volume were: 57, 65, 128, 336, 133, 83 and 41. Panel A displays mean childhood IQ (average from measurements at 7, 9 and 11) for each of the WMH volume groups. Panel B displays the mean adult IQ (measured at age 45) for the same groups. Panel C displays the association between WMH volume and cognitive decline. The average IQ in childhood and adulthood in each of these is plotted to illustrate increasing severity of cognitive decline as WMH volume increases. All error bars display the standard error of the mean.

Secondary analyses using the four subscales of IQ showed that larger WMH volume was associated with lower scores on processing speed (ß = −0.14, 95% CI = −0.20 to −0.07; p < 0.001), verbal comprehension (ß = −0.14, 95% CI = −0.21 to −0.07; p < 0.001), and perceptual reasoning (ß = −0.13, 95% CI = −0.19 to −0.06; p < 0.001). There was also a trending association between larger volume and decreased working memory ability (ß = −0.06, 95% CI = −0.13 to 0.00; p = 0.06).

## Discussion

In a population-representative birth cohort of individuals now in midlife, we found that WMHs are 1) common, 2) associated with cognitive abilities in childhood and midlife, and 3) associated with cognitive decline from childhood to midlife. While there is robust evidence that WMHs are related to cognitive decline in older adults (Debette and Markus, 2010; Prins and Scheltens, 2015), our study provides initial evidence that this relationship begins by midlife.

These findings demonstrate that the link between WMHs and early signs of cognitive decline are detectable decades before clinical symptoms of ADRD typically emerge. Longitudinal studies have shown that WMHs tend to grow and expand from existing lesions and that higher baseline volumes predict faster accumulation of WMHs and more rapid cognitive decline in older adults (Maillard *et al.*, 2012; Prins and Scheltens, 2015). Thus, our findings suggest that WMHs may be a surrogate biomarker for identifying individuals in midlife who are at risk for future clinically significant cognitive decline or ADRD. Our results further show that accumulation of WMHs in midlife already indicates mild cognitive decline. This is important because even subclinical cognitive decline impacts daily functioning and psychological well-being (Tucker-Drob, 2011).

Interestingly, our results also showed a modest association between low childhood IQ and WMHs in midlife. This finding suggests at least two potential pathways for the development of WMHs. The first possibility is that children with lower IQs tend to be born into or seek out environments that lead to higher rates of neurodegeneration (e.g., poor nutrition, smoking, drug abuse, lead exposure). Over time these exposures may lead to negative health outcomes, such as higher risk for cardiovascular disease, brain damage, and higher blood pressure, which contribute to increased WMH volume in midlife. This perspective would suggest that interventions to limit neurodegenerative environmental exposures (e.g., anti-drug messaging, better nutrition) in high-risk children could limit the burden of cognitive decline and ADRD later in life. The second possibility is that low IQ is an indicator of lower overall brain integrity that was present early in life (Deary, 2012). This perspective suggests that the association between low childhood IQ and midlife WMH is driven by a higher vulnerability to tissue damage and faster neurodegeneration in low-IQ children, given the same lifetime exposures. This further suggests a need for interventions that increase brain resiliency and boost tissue regeneration in those at highest risk (e.g., cognitive training or pharmaceutical intervention). A limitation of our study is the lack of childhood neuroimaging to assess the development of WMHs across the lifespan, although it should be noted that no sample with WMH measures in midlife would have childhood WMH measures, because cohorts of non-patient children did not have MRI imaging 40 years ago. As such, our findings point to the need to investigate these possible mechanistic pathways in future studies with child-to-adult imaging data.

Intervention efforts targeting WMHs as a surrogate biomarker in older adults have had mixed results (Prins and Scheltens, 2015). One reason for this inconsistency could be that older adults have accumulated decades of irrevocable age-related tissue damage. Given that prevention of damage is often more efficacious than reversal of damage (Sperling *et al.*, 2014; Moffitt *et al.*, 2017), particularly in the brain, our results suggest that lifestyle and pharmaceutical interventions aimed at slowing the progression of WMHs in midlife may be promising complements to interventions in older adults. Due to their compounding growth during aging, WMHs may be especially useful for selecting individuals in midlife who are at highest risk for future cognitive decline and who may most benefit from early prevention.

## Acknowledgements

Thank you to members of the Advisory Board for the Dunedin Neuroimaging Study, the Dunedin Study members, Unit research staff, and Study founder Phil Silva.

## Funding

This research was supported by National Institute on Aging grants R01AG032282, R01AG049789, and UK Medical Research Council grant MR/P005918. Additional support was provided by the Jacobs Foundation. MLE is supported by the National Science Foundation Graduate Research Fellowship under Grant No. NSF DGE-1644868. The Dunedin Multidisciplinary Health and Development Study is supported by the NZ HRC and NZ MBIE.

### Competing interests

The authors declare no competing interests.

