## Supplemental Info for "White matter hyperintensities are common in midlife and already associated with cognitive decline"

**Supplementary Material**

*Phase 45 Attrition Analysis*

We conducted an attrition analysis using childhood intelligence quotient (IQ; Supplementary Fig.1) and socioeconomic status (SES; Supplementary Fig. 2) to determine whether participants in the Phase 45 data collection were representative of the original cohort.

**Attrition Analysis of Childhood IQ in Phase 45**

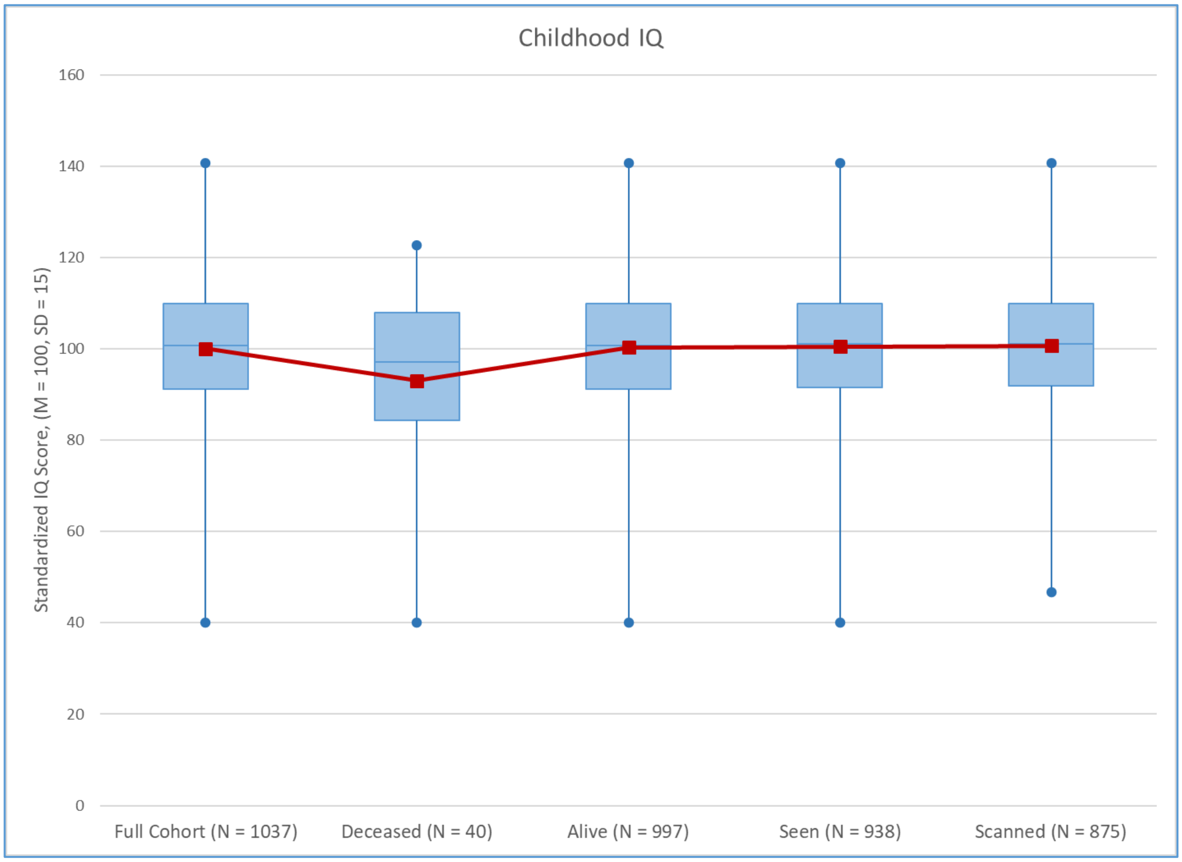


**Supplementary Figure 1.** No significant differences in childhood IQ were found between the full cohort, those still alive, those seen at Phase 45 or those scanned at Phase 45. Those who were deceased by the Phase 45 data collection had significantly lower childhood IQ's than those who were still alive ($t = 2.09$, $p = 0.04$).

**Attrition Analysis of Childhood SES in Phase 45**

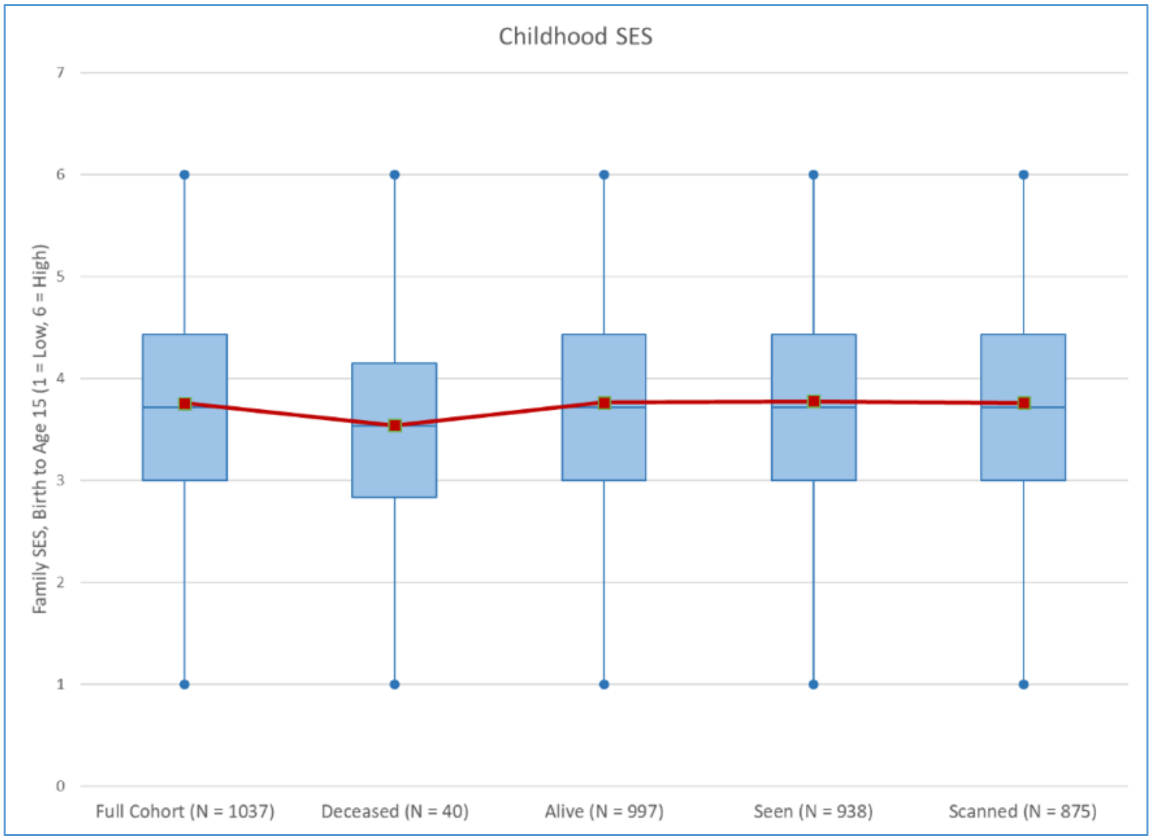


**Supplementary Figure 2**. No significant differences were found between the full cohort, those deceased, those alive, those seen at Phase 45 or those scanned at Phase 45 on childhood SES.
